# Sensitive detection of rare disease-associated cell subsets via representation learning

**DOI:** 10.1101/046508

**Authors:** Eirini Arvaniti, Manfred Claassen

## Abstract

Rare cell populations play a pivotal role in the initiation and progression of diseases like cancer. However, the identification of such subpopulations remains a difficult task. This work describes CellCnn, a representation learning approach to detect rare cell subsets associated with disease using high dimensional single cell measurements. Using CellCnn, we identify paracrine signaling and AIDS onset associated cell subsets in peripheral blood, and minimal residual disease associated populations in leukemia with frequencies as low as 0.005%.

## Main

Health and disease status of multicellular organisms pivotally depends on rare cell populations, as for instance hematopoietic stem cells or tumor initiating cell subsets^1^. Advances in single cell-resolved molecular measurement technologies have increasingly enabled us to describe cell population heterogeneity and, specifically, rare subpopulations in health and disease^2^. It is becoming routine to measure thousands of DNA, RNA^3^ and dozens of protein^4^ species in more than thousands of single cells, optionally including their spatial context^5–7^.

Such multiparametric single cell snapshots have been used to define heterogeneous cell population structure using unsupervised clustering techniques that generate a *representation* of a cell population, defined in terms of cluster-based features such as cluster medians^8^. While unsupervised machine learning constitutes a powerful exploratory tool, the identification of disease-associated cell subsets requires a further supervised learning step to associate the clustering derived representation with disease status. Unsupervised approaches have been extended to the classification of single cell samples and have been successful where disease association manifested itself in condition-specific differences of abundant cell subpopulations^8,9^.

Unsupervised approaches describe general population features that are not necessarily associated with disease status. Typically a large number of cell population features (i.e. thousands^9^ or millions^10^) are required to detect rare cell subsets from high dimensional measurements (i.e. 20+ dimensions). Most such features are not relevant, leading to overfitting or even precluding the identification of disease-associated rare cell populations. As this study will demonstrate, this situation severely limits the capacity of existing approaches to take advantage of novel highly multiparametric single cell measurements to yield insights into the disease mechanisms such as minimal residual disease or tumor-initiating cells^1^.

CellCnn overcomes this critical limitation and facilitates the detection of rare disease-associated cell subsets. Unlike previous methods, CellCnn does not separate the steps of extracting a cell population representation and associating it with disease status. Combining these two tasks requires an approach that (1) is capable of operating on the basis of a set of unordered single cell measurements, (2) specifically learns representations of single cell measurements that are associated with the considered phenotype and (3) takes advantage of the possibly large number of such recordings. We bring together concepts from *multiple instance learning* and *convolutional neural networks*^11^ to meet these requirements. CellCnn takes as input a set of biological specimens (multi-cell inputs) that are each associated with a phenotype. Such biological specimens and phenotypes comprise, for instance, patient blood or tissue samples with their associated disease status or survival information. It is difficult to automatically learn the molecular basis of this association since it possibly manifests itself by differences of an, a priori unknown, cell subset. To address this difficulty, CellCnn associates a multi-cell input with the considered phenotype by means of a convolutional neural network. The network automatically learns a concise cell population representation in terms of molecular profiles (filters) of individual cells whose presence or frequency is associated with a phenotype (**Fig. 1a, Methods**). We applied CellCnn (1) in a classification setting to reconstruct cell type specific signaling responses in samples of peripheral blood mononuclear cells, (2) in a regression setting to identify abundant cell populations associated with disease onset after HIV infection, and achieved comparable prediction accuracy to a state of the art analysis performed recently^9^, however with computational cost reduced by several orders of magnitude. Finally, we considered the task of minimal residual disease detection in leukemia to demonstrate the unique ability of CellCnn to identify extremely rare phenotype-associated cell subsets.

**Figure 1.**
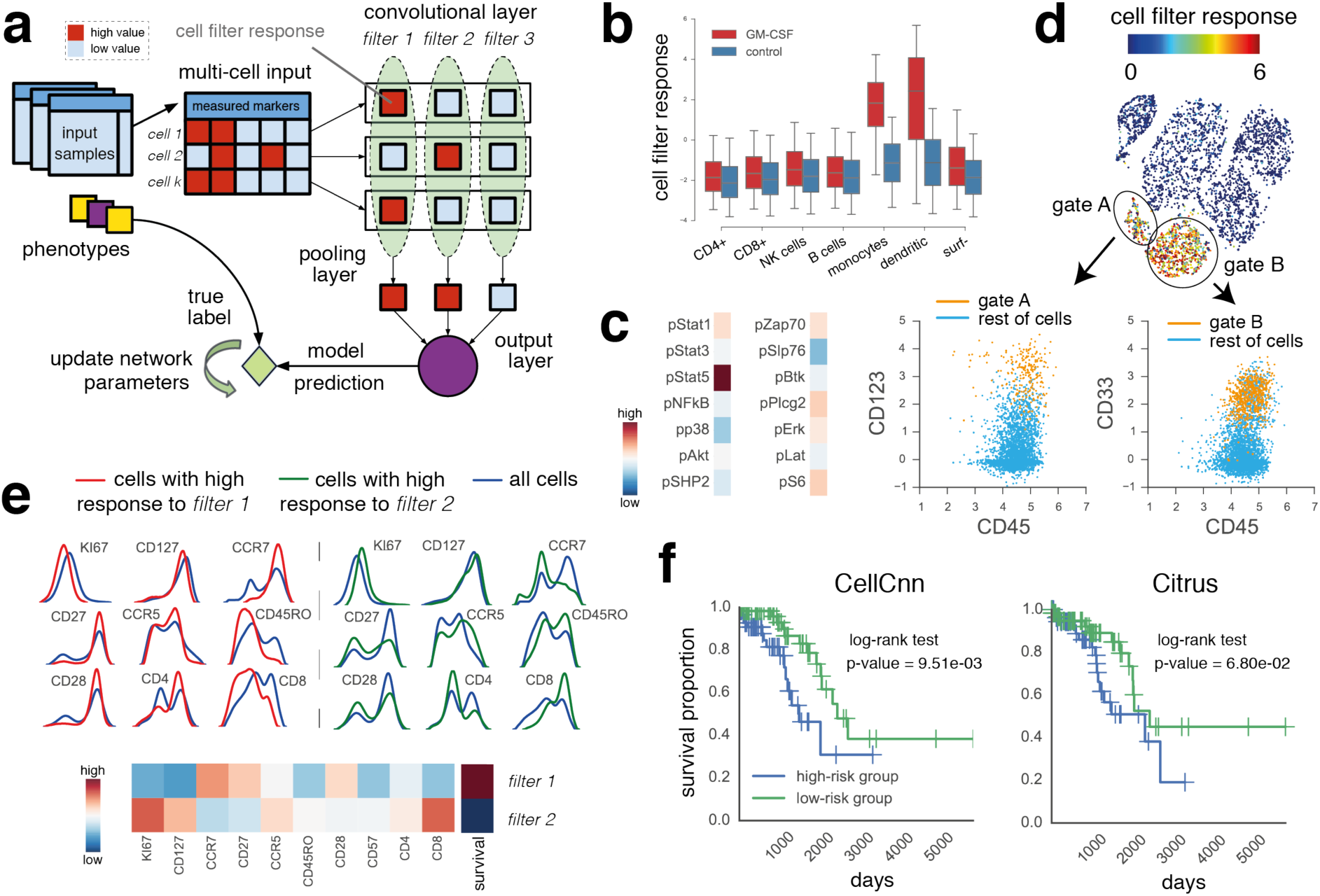
CellCnn overview and demonstration. **(a)** CellCnn convolutional neural network architecture. CellCnn takes as input the single cell measurements (multi-cell input), each of which is annotated with a phenotype. Node activities in the convolutional layer are defined as weighted sums over single cell molecular profiles. Nodes in the pooling layer evaluate the presence (max pooling) or frequency (mean pooling) of specific cell subsets. The output of the network estimates the sample-associated phenotype (e.g. disease condition, expected survival). Network training optimizes weights to match training dataset phenotype. Trained filter weights correspond to molecular profiles of relevant cell subsets and allow for assignment of the cell subset membership of individual cells (cell filter response). **(b)** CellCnn classification of GM-CSF (un-) stimulated peripheral blood mononuclear cell populations monitored with mass cytometry. Response of individual cells (grouped by manually gated cell types) is shown for both conditions. Significantly higher cell filter response for monocytes and dendritic cells in the stimulated sample. **(c)** Filter weights learned for GM-CSF stimulus. **(d)** t-SNE projection using all cell type-defining surface markers **not** used by CellCnn, coloured by cell filter response. High cell filter response regions (**Supplementary Fig. 7**) map to monocytes (CD33+) and dendritic cells (CD123+). **(e)** Reconstruction of cell subsets predicting AIDS-free survival in HIV-infected patients. (**f**) Kaplan-Meier plots for high and low risk patient cohort according to CellCnn survival prediction (p=9.51e-03, log-rank test, computation time: 5min, single laptop core) and state of the art: Citrus (p=6.8e-02, 3 days, 24 Intel Xeon cores).

First, we applied CellCnn to a mass cytometry dataset acquired from samples of peripheral blood mononuclear cells^12^. These samples were exposed to various paracrine agents and their proteomic responses recorded at the single cell level with respect to 14 intracellular markers and 11 cell-surface markers characteristic of immune cell type. CellCnn was trained for each paracrine agent to classify stimulated and unstimulated samples using only the 14 intracellular markers. CellCnn learned two filters that were used to compute the weighted sum of the abundance profile for each single cell (cell filter responses, **Fig. 1a**). We investigated the cell type-specific filter responses and found very specific and sensitive enrichment of the cell types expected to specifically respond to the considered agent, i.e. differential response by monocytes and dendritic cells in the case of GM-CSF exposure^13^ (**Fig. 1b/d**, see **Supplementary Fig. 1** for the remaining agents considered in^12^). The learned filter positively associated with GM-CSF stimulation is, as to be expected, predominantly activated by cells with high pStat5 levels (**Fig. 1c**, see **Supplementary Fig. 2** for filters learned for the remaining agents).

We used CellCnn to identify T cell subsets associated with increased risk of AIDS onset in a 383 HIV-infected patient cohort^14^. Flow cytometry measurements of 10 T cell related molecular markers from peripheral blood, and AIDS-free survival time were available for each individual. Trained on a subcohort of 256 individuals, CellCnn identified cell subsets with either elevated proliferation marker Ki67, or naive T-cell phenotype, whose frequency has been reported to be associated with AIDS-free survival (**Fig. 1e**)^15^. CellCnn was further used to categorize the remaining set of 127 test individuals into a low- and high-risk group (**Methods**). Kaplan-Meier curves of these groups are significantly different (p-value = 9.51e-03, log-rank test, **Fig. 1f**). Citrus, a state of the art approach to identifying clinically prognostic cell subsets^9^ achieved a less significant dissection of both groups on the same training and test data partition (p-value = 6.8e-02, **Fig. 1f**).

We assessed the ability of CellCnn to detect rare cell populations associated with minimal residual disease in acute lymphoblastic leukemia (ALL) and acute myeloid leukemia (AML). Specifically, we analyzed mass cytometry datasets of healthy bone marrow samples with leukemic blast spike-in subpopulations of decreasing frequency to mimic the minimal residual disease (MRD) phenotype^16,17^. CellCnn correctly identified the ALL and AML MRD associated cell subset at a frequency as low as 0.4/0.1 % (500/720 blast cells) respectively. We found CD45, CD34, CD10 and respectively CD47, CD7, CD44, CD38 differentially expressed in the predictive subsets, matching the expectations for MRD in ALL (**Fig. 2a**) and AML (**Fig. 2d**). CellCnn achieved an almost perfectly precise recovery of the MRD associated cell subset for a recall of 80% (**Fig. 2b/e**). We compared CellCnn with (1) a state-of-the-art distance-based outlier detection algorithm^18^, constituting a quantifiable variant of visually inspecting condition specific projection map differences (e.g. t-SNE maps^16,19^), (2) logistic regression classifiers that take as input either single cell profiles or the average multi-cell input profiles and (3) Citrus^9^, all failing to identify the ALL/AML MRD subsets with reasonable precision (**Fig. 2b/e, Supplementary Fig. 3**, details see **Methods**). We further considered more extreme situations with decreasing frequency of the MRD associated cell subset down to 0.08/0.005 % (100/38 blast cells) for ALL/AML (**Fig. 2c/f**). While the task of recovering the correct cell subset becomes increasingly difficult (**Supplementary Fig. 4**), cell classification precision of CellCnn stayed considerably high for all considered MRD cell subset frequencies.

**Figure 2.**
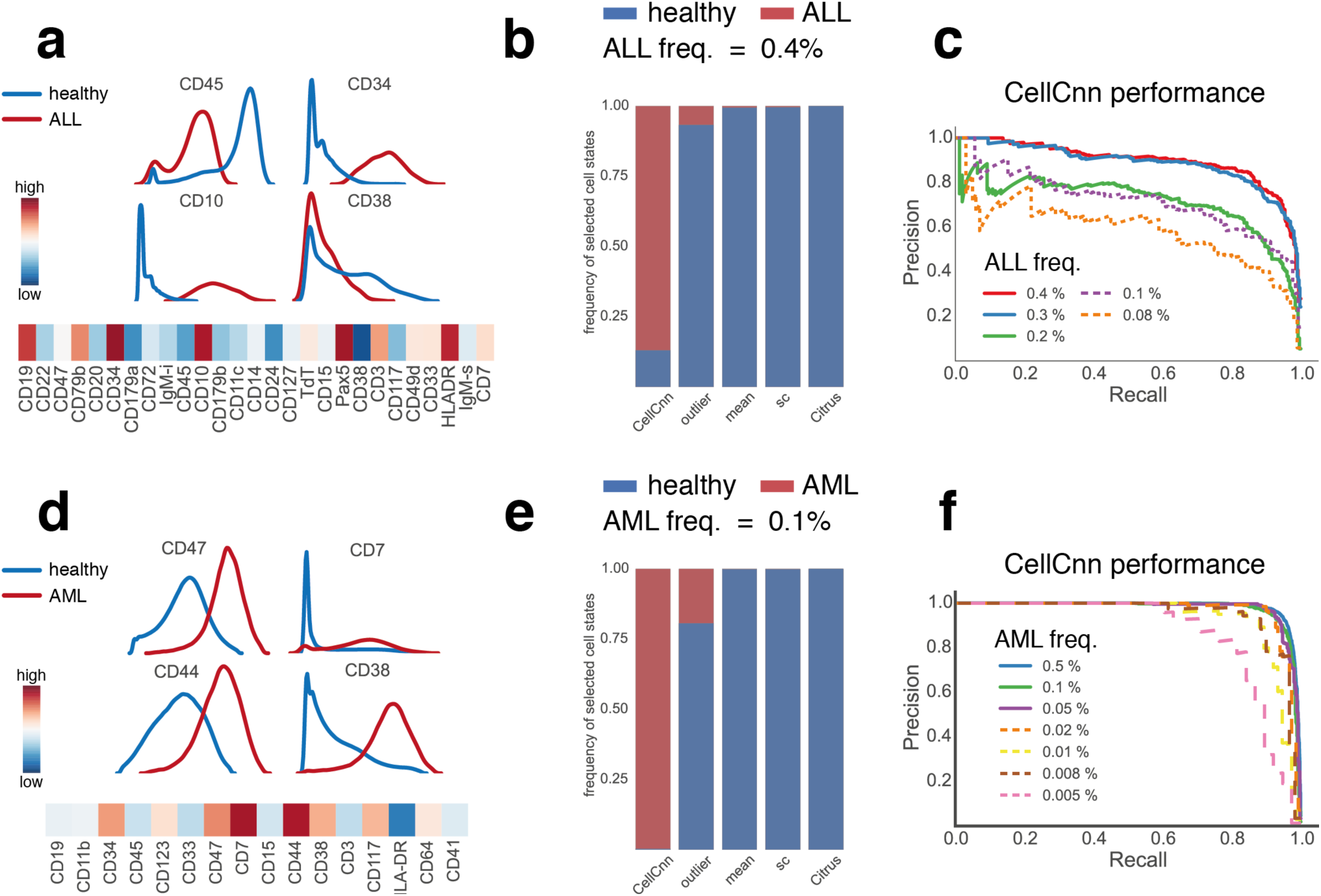
Identification of rare cell populations associated with minimal residual disease in acute lymphoblastic (ALL) and acute myeloid leukemia (AML). (**a**) Marker histograms of healthy and ALL populations (ground truth) and ALL-specific filter weights learned by CellCnn. (**b**) Comparison to baseline methods (outlier: distance-based outlier detection^18^, mean: logistic regression on multi-cell input summary profiles, sc: logistic regression on single cell profiles, Citrus^9^) for ALL MRD population at 0.4%. For the first four methods, we report the disease condition of top-scoring cells for a recall of 0.8. Citrus does not provide a precision-recall series, therefore cells assigned to clusters with differentially abundant markers are reported. (**c**) Precision-recall of CellCnn for various ALL MRD cell population frequencies. Solid lines indicate that an automated procedure (described in **Methods**) was used for selecting the reported filter. Dashed lines indicate that the filter achieving the highest area under the precision recall curve (AUPRC) score is reported (details in **Methods**). (**d**) Marker histograms of healthy and AML populations (ground truth) and AML-specific filter weights learned by CellCnn. (**e**) Comparison to baseline methods for AML MRD population at 0.1%. (**f**) Precision-recall of CellCnn for various AML MRD cell population frequencies. Solid and dashed lines are interpreted as in (c).

CellCnn achieves this unprecedented high precision by overcoming the inherent limitations of the unsupervised feature engineering strategies of state of the art approaches. When analyzing samples from modern single-cell techniques with increasing multiparametricity, such as mass cytometry, these approaches enumerate tens of thousands^9^ or an exponential number^10^ of features, at the cost of accumulating many potentially uninformative, confounding features. This situation leads to both computational bottlenecks and loss of statistical power (**Fig. 1f**). CellCnn provides a solution to this limitation by jointly and thereby efficiently solving the feature engineering, selection and association tasks in a single supervised learning step. Further, CellCnn training is efficient, scaling linearly with the number of measured components. Consequently, CellCnn is applicable to a variety of highly multiparametric single cell data sources beyond flow and mass cytometry, such as single cell RNA sequencing or imaging data.

CellCnn is expected to enable the discovery of new disease-associated cell populations such as tumor initiating cells from patient cohorts of suitable size or longitudinal studies. These results lend themselves to the design of personalized diagnosis and treatment modalities. Given the expected increase in patient cohort sizes in concerted initiatives as the The Cancer Genome Atlas (TCGA) and concomitant rise in their analysis with single cell technologies^20^, we expect scalable representation learning approaches such as CellCnn to uniquely take advantage of the resulting data by enabling the discovery of disease mechanisms mediated by rare cell populations, in both basic research and personalized medicine.

## METHODS

### Datasets

The mass cytometry data set of peripheral blood mononuclear cells (PBMC) and the flow cytometry dataset of HIV infected patients were adopted respectively from Bodenmiller *et al*.^12^ and the U.S. Military HIV Natural History Study^14^.

The mass cytometry datasets for the MRD study were based on the ALL samples from Amir *et al*. *^16^* and the AML samples from Levine *et al*.^17^. Specifically, in the study from Amir *et al*. metal-barcoded cells from an ALL patient sample were spiked into a healthy bone marrow sample. The abundance of the leukemic population was 0.4% of the total cell counts in the mixed sample. We created additional datasets with decreasing frequencies by selecting random subsets of the leukemic population with frequencies 0.3%, 0.2%, 0.1% and 0.08%. In each case, the mixed sample is compared with a control sample from healthy bone marrow of a different patient.

Also, we have created synthetic AML MRD samples by computationally combining AML CD34+ gated blast cells (Supplementary Fig. 8) and cells from a healthy bone marrow into mixed synthetic samples on the basis of the data provided by Levine *et al*. Different mixed samples have been created by introducing different numbers of blast cells from different AML patients 17. The considered frequencies of AML blasts are 0.5%, 0.1%, 0.05%, 0.02%, 0.01%, 0.008% and 0.005%. In each case, the mixed sample is compared with a pool of control samples from four different healthy bone marrows.

### CellCnn network architecture and training

CellCnn aims at identification of disease-associated cell subpopulations, which is an example of a *multiple instance learning* task^21^. Such learning tasks consist of learning the relation between labels (e.g. disease status) and data items that manifest themselves as bags (multi-cell inputs) of instances (cell molecular profiles). We address this multiple instance learning task with a convolutional neural network approach. CellCnn implements a variant of a convolutional neural network. Such networks are artificial neural networks originally designed to process the two-dimensional structure of images and typically consist of one or more sets of convolutional and pooling layers^11^. Briefly, the convolutional layer comprises filters that evaluate the occurrence of specific patterns in image patches and the pooling layer computes summaries of these occurrences. We adapted the convolutional neural network architecture to process unordered multi-cell inputs. Image patches correspond to individual cell measurements. Each cell measurement was evaluated with respect to every convolutional filter, i.e. to its fit to respective molecular profile (cell filter response) in the convolutional layer. The computation at the pooling layer consisted of either selecting the maximum (max-pooling) or mean (mean-pooling) response within the multi-cell inout^11^. Pooling was performed separately for each convolutional filter. Max-pooling computes the maximum response over all members of a multi-cell input for a particular filter, and thereby measures the presence of cells yielding high cell filter response. Max-pooling was performed for the analysis of the peripheral blood, ALL and AML datasets, where cell presence appeared to be most informative. Mean-pooling evaluates the average cell filter response of a multi-cell input, and thereby serves as an approximation of the frequency of the cell subset strongly responding to a specific filter. Cell subset frequencies turned out to be most informative for the analysis of the HIV dataset and therefore mean-pooling was performed. Finally, the pooling layer was connected to the output layer. For regression problems the output layer contains a single node, whereas for classification problems it contains one node per class. Nodes in the output layer compute a weighted sum over the pooling layer nodes, followed by a nonlinear operation (hyperbolic tangent for regression and softmax for classification, further details in Supplementary Notes).

The convolutional filter weights and output layer weights were optimized for optimal association of multi-cell inputs with their phenotype labels using mini-batch stochastic gradient descent with Nesterov momentum^22^. We adopted a set of commonly used hyperparameter settings (`learning rate = 0.03, momentum = 0.9, mini-batch size = 128, maximum number of training epochs = 20`) and kept them fixed in all our experiments. Two or three convolutional filters were used to encourage model simplicity and avoid overfitting. Network weights were initialized randomly from a uniform distribution. We minimized the multinomial logistic regression objective for classification and mean squared error for regression, both augmented with small L^2^ weight decay in the order of 10^-8^ (more details on the CellCnn methodology in Supplementary Notes).

### Cell subsets as multi-cell inputs

To take advantage of high content single cell techniques like flow or mass cytometry, CellCnn optionally takes multiple random cell subsets of a specific cytometry sample as input to increase the effective number of data points for association.

In all our experiments, random cell subsets, drawn with replacement from the original cytometry samples, were used as multi-cell input training examples of CellCnn. Multi-cell inputs were chosen sufficiently large, to ensure that these contain cells with the molecular profile of interest according to the expected frequency. From our experiments on mass cytometry data, we found that generating 4096 multi-cell inputs per class, with each multi-cell input comprising 1000 cells, performed well (Supplementary Fig. 5). If we are interested in extremely rare populations (abundance < 1%) then we use a modified procedure for creating multi-cell inputs. 50% of a multi-cell input is sampled uniformly at random from the whole cell population whereas the other 50% is sampled from cells with high outlierness score. We define the outlierness score of each cell based on the distances between this cell and its closest neighbors from the control samples 18 (details in Supplementary Notes).

### Model selection and interpretation

#### PBMC and ALL/AML datasets

Each sample was initially split into a training (80%) and a validation (20%) set of cells. We trained 20 models with uniformly random initial weights and pre-selected the models with predictive accuracy higher than 99% on the validation set. Then we performed a type of stability selection procedure to prioritize frequently occurring filters. Hierarchical clustering with cosine similarity as metric was performed on the matrix of filter weights and flat clusters were formed by cutting the dendrogram at a cosine similarity of 0.7. From each cluster, the filter with minimum sum of distances to all other members of the cluster was chosen as representative. Finally, the filter representative of the biggest cluster with positive weight connection to the output node corresponding to stimulation/leukemia was chosen for the detection of stimulated/diseased cells.

The automated filter selection procedure described above was not performed when the leukemic blast population comprised less than 200 cells. In those cases it was likely that leukemic cells might not be included in the validation set and thus the purpose of validation would be lost. In the absence of a trustworthy validation set, we were not able to prioritize specific filters. Therefore, we examined if at least one trained filter was able to detect the rare leukemic population.

#### HIV-cohort dataset

The full patient cohort was randomly split into a training (2/3) and a test (1/3) cohort. We used 10-fold cross validation on the training cohort resulting in 10 models, each trained on a different subset of the training cohort. Information from all cross validation runs was used to select frequently occurring filters. We compiled a matrix of all filter weights from the 10 networks and performed hierarchical clustering using cosine similarity as metric. Clusters were determined by cutting the dendrogram at a cosine similarity of 0.7. From each cluster with at least 5 members, the cluster centroid was chosen as representative. Representative filters are depicted in Figure 1e. Finally, an ensemble model, consisting of the top 5 networks with best validation predictive performance from the 10 cross validation runs, was used to predict survival times for the individuals in the test cohort. For the test phase, one subset of 3000 cells was used per individual. The output of CellCnn corresponded to predicted disease-free survival time for each patient and was used to split the test cohort into low- and high-risk groups. The threshold used for defining the two risk groups was the median predicted survival time.

### Baseline cell population models

The following models were used for comparison with CellCnn.

Outlier detection: We used a state of the art distance-based outlier detection method^18^. A set S of s observations (single cell profiles) is randomly sampled from the *inlier* class (i.e. the healthy control samples) and then used to evaluate the *outlierness* of single cell profiles in the test samples. The outlierness of an observation is defined as the L_1_ distance between this observation and its closest neighbour in S. Results for different values of s are given in Supplementary Figure 6. We finally used s = 200,000.

Single-cell input logistic regression: a logistic regression classifier that takes as input single cell profiles. Each single cell profile is labeled with the label (e.g. disease condition, survival time) of its corresponding cytometry sample.

Multi-cell input logistic regression: a logistic regression classifier that operates on the same multi-cell inputs as CellCnn. Mean abundances of multi-cell input serve as features for the classifier. Each multi-cell input is labeled with the label of its corresponding cytometry sample.

Citrus: a state-of-the-art approach for detecting phenotype-associated cell subpopulations^9^. Citrus initially performs hierarchical clustering of single cell profiles from all considered cytometry samples, selects the clusters that contain at least a minimum number of cell events (according to the minimum cluster size threshold that is defined by the user) and computes cluster-based features (e.g. population medians or abundances) individually for each cytometry sample. The computed cluster-based features are used as input to a L_1_ -regularized predictor that detects phenotype-associated differentially abundant features. See Supplementary Notes for a detailed description of the parameters used in individual Citrus runs.

### Code availability

CellCnn is implemented in Python 2.7 and uses the neural network libraries Theano^23^ and Lasagne/Nolearn^24^. It is available for download at https://github.com/eiriniar/CellCnn.

## Acknowledgements

EA is funded by the SystemsX.ch RTD project PhosphonetPPM. We thank Justin Feigelman, Will Macnair and Uwe Sauer for helpful comments on the manuscript.

## Author contributions

M.C. and E.A. conceived the CellCnn methodology, and designed experiments. E.A. implemented the CellCnn method and carried out the experiments. M.C. and E.A. wrote the manuscript.

## Correspondence

Correspondence should be addressed to M.C.

